# Brain Control of a Computer Cursor for Online Target Selection - A Non-Invasive BCI for Continuous Movement Decoding

**DOI:** 10.64898/2026.06.23.733968

**Authors:** Markus R. Crell, Kyriaki Kostoglou, Patrick Suwandjieff, Johanna Egger, Gernot R. Müller-Putz

## Abstract

Non-invasive brain-computer interfaces (BCIs) have substantially advanced in the field of continuous cursor control over the past decade. Yet, current methods lack key control aspects such as initiation and termination of cursor movements as well as evaluation in real-world applications. In this study, we introduce a framework for continuous, electroencephalography-based cursor control that supports both active movement and no-movement states, thereby allowing for inactive periods of the user when no control input is desired. We demonstrate its applicability in healthy participants and show its performance in real-world application through the selection of targets on a screen. This demonstrates that participants can leverage the continuous control cursor control and the intentional starting and stopping of motions to effectively select targets on a screen through dwell-time selection. On average, 7.1 out of 40 targets were correctly selected (level of significant performance: 4.5 targets), while experienced BCI users achieved an average of 12.8 targets. The proposed framework additionally demonstrates compatibility with motor-impaired people without residual hand motions since it does not rely on observable movements for model training.

## 1 INTRODUCTION

Continuous cursor control is a prerequisite for intuitive human-computer interaction. Yet, most non-invasive brain-computer interface (BCI) systems emphasize discrete classification rather than continuous motion. For severely motor-impaired patients, such as patients with Locked-in Syndrome (LIS), the control of a continuous input method is especially valuable since these patients rely on caregivers or eye tracking systems as their only form of communication. Current invasive BCIs have substantially progressed in this area and demonstrate remarkable continuous control of computer cursors (Wilson et al., 2025; Deo et al., 2024; Dekleva et al., 2021; Degenhart et al., 2018).

However, invasive methods require complex surgical interventions which are associated with medical risks and may thus not be feasible for all patients. They also require a lengthy planning and screening phase during which participants do not benefit from the BCI. Non-invasive BCIs can offer a practical alternative. Due to their straightforward and quick application, they can be made available to nearly all patients at a fraction of the cost and effort of invasive BCIs. However, non-invasive BCIs utilizing electroencephalography (EEG) deal with a low signal-to-noise ratio and oftentimes poor performances in distinguishing fine motions of the hand or other body parts (Edelman et al., 2024). While the online classification of discrete (hand) movements versus rest and the utilization in matrix spellers can be achieved with high accuracies and low rates of false positives per minute (Suwandjieff et al., 2026; Crell et al., 2026; Ibanez et al., 2014; Xu et al., 2014; Fatourechi et al., 2007; Scherer et al., 2004; Pereira et al., 2021), reliable EEG-based cursor control remains an unsolved problem.

Prior work on this topic has largely followed two approaches: (i) the direct decoding of continuous kinematics and (ii) high-rate classification of discrete movements. In the first approach, continuous kinematic variables such as position, velocity, or direction are regressed directly from EEG. This approach facilitates closed-loop, intuitive trajectory control by mapping attempted or executed movement to kinematics. Prior studies utilizing this approach have demonstrated online cursor control by directly regressing hand position in both able-bodied and motor-impaired participants (Mondini et al., 2024, 2020; Pulferer et al., 2022). Without online application, other studies further showed promising methods for the continuous regression of speed, movement direction and the simultaneous decoding of both (Lu et al., 2026; Crell and Müller-Putz, 2024b; Kobler et al., 2020b) from hand motions. The second approach detects discrete gestures at high temporal resolution and converts them into directional commands. Although this pseudo-continuous strategy lacks the natural, intuitive control associated with direct regression of continuous movements from a single end-effector, it was proven to enable directional control of virtual cursors in participants (Edelman et al., 2019; Suma et al., 2020; Forenzo et al., 2024).

While both approaches demonstrate the feasibility of generating continuous trajectories from non-invasive EEG, evaluation is largely restricted to pursuit tracking tasks, with performance primarily quantified using correlation coefficients or trajectory error metrics. Importantly, none of the current studies systematically assess real-world cursor control applications such as target selections, leaving the practical applicability of continuous EEG-based cursor control largely insufficiently established. Moreover, current continuous and pseudo-continuous EEG cursor control studies do not explicitly define or evaluate a robust stopping or no-movement state. Cursor motion is typically updated continuously, requiring sustained neural modulation to suppress movement rather than enabling deliberate idle periods (Wolpaw and McFarland, 2004). The absence of a dedicated stopping mechanism poses a critical barrier for real-world use, where stable holding, target dwelling, and intentional inactivity are essential. While some previous work has successfully identified start and stop of continuous movements by detecting movement on- and offsets (Crell and Müller-Putz, 2024a), an extension to continuous cursor control has not been attempted so far.

To address these limitations, our study proposes a continuous cursor control framework that enables explicit no-movement periods of the cursor while preserving adequate directional control during movement periods. We further demonstrate, for the first time in EEG-based BCIs, successful target selection with explicit no-movement periods using continuous EEG decoding.

## 2 RESULTS

### 2.1 Online Cursor Control for Target Selection

Fifteen participants controlled a cursor on a screen using the speed (binary, movement vs. rest) and direction (velocity in x- and y-direction) of their hand movements decoded from EEG (Figure 1 A). The cursor control was evaluated during 40 trials of an online target selection task (Figure 1 B). On average, 7.1 ± 6.1 (17.7 ± 15.3 %) targets were correctly selected during the online task with an average time of 18.01 s until selection (including 3.0 s of dwell-time for selection). To evaluate whether the observed performance may have occurred by chance, we tested the null hypothesis that the probability of selecting a target was independent of its position, i.e., the executed trajectories were unrelated to the targets (Crell and Müller-Putz, 2026). The resulting level of significant performance (p*_s_*) was exceeded by the actual target-selection performance of the participants in 8 cases. 7 participants did not achieve a performance better than p*_s_* and thus did not achieve a cursor control that was significantly better than random. On average, the level of significant performance was p*_s_* = 4.5 target selections.

**Figure 1.**
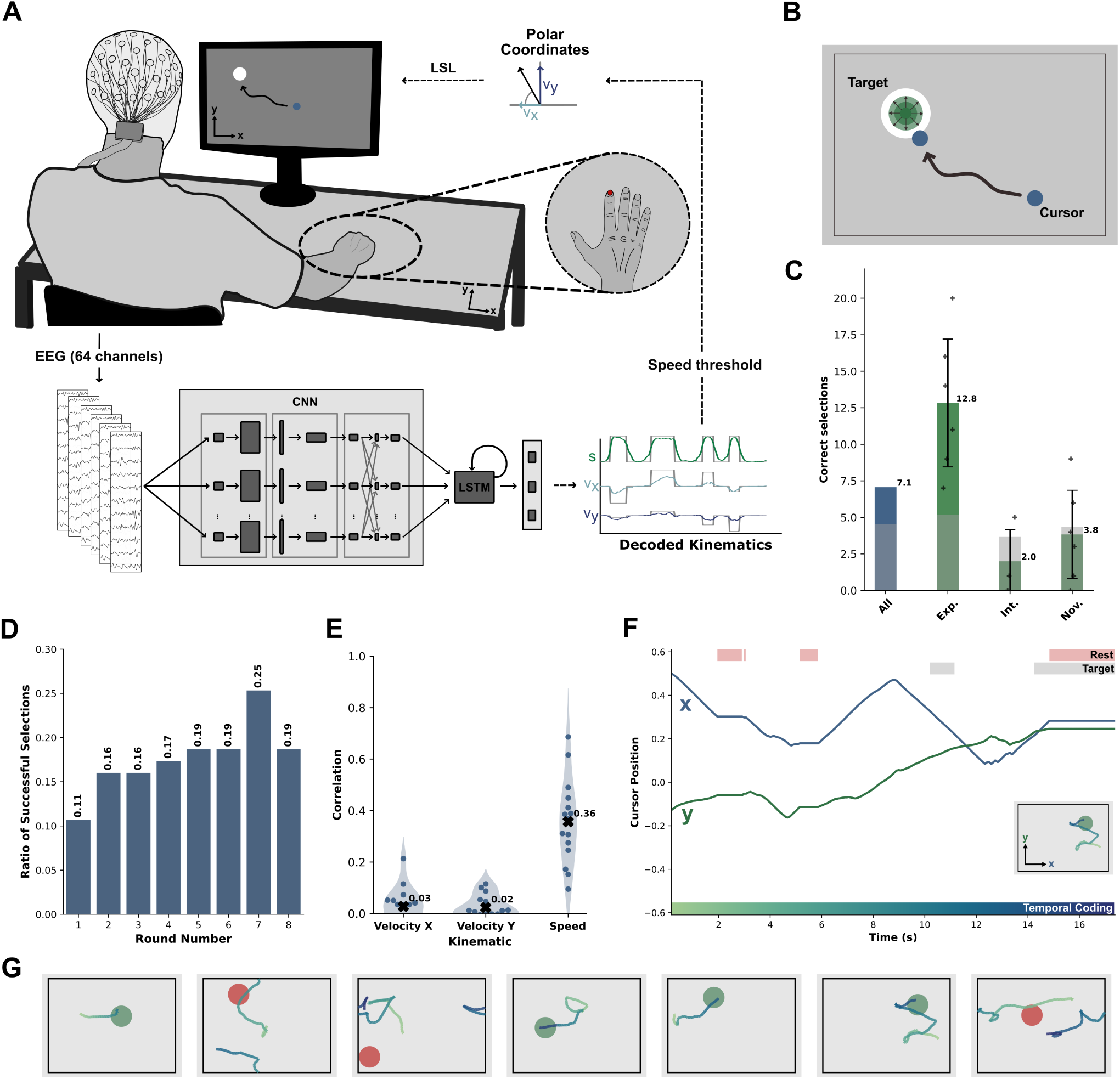
Overview of the online cursor control framework. (A) BCI setup of the online cursor control with data processing pipeline and cursor control algorithm. (B) Online cursor control task including moving the cursor and selecting the target. (C) The overall number of selected targets and the performance grouped by prior participant experience with BCI systems. The grey area indicates the average level of selected targets above which a performance can be considered significantly better than chance. (D) Grand average ratio of correctly selected targets per round. (E) Participant-wise correlation between the executed and the decoded velocities in x- and y-direction and the speed. (F) Continuous cursor position over time during an online target selection trial. Rest (no-movement) is coded in red and time spent inside the target is coded in grey. (G) Examples of movements executed during the online target selection task. Correctly selected targets are shown in green, while unselected targets are shown in red. The temporal aspect of the motion is coded according to the temporal coding scheme shown in panel F.

We further identified trends over time (training effects) for the target selection performance and time to select a target. Each additional round was associated with higher odds of success (odds ratio per round = 1.10, 95 % CI: 1.03 − 1.16, p = 0.003), showing that the ratio of correctly selected targets per round increased significantly over time. Figure 1 D shows the number of selected targets per round over time, visibly demonstrating the learning effect. We did not find any significant change in the time required to correctly select a target.

We additionally assessed the decoder performance for decoding the speed and direction of the actual movement (Figure 1 E). The Pearson correlation coefficient between the executed motion and the decoded movement during the online target selection task was 0.36 ± 0.16 for the speed and 0.03 ± 0.07 and 0.02 ± 0.05 for the velocities in x- and y-direction, respectively. The accuracy of the speed detection was extracted by applying a threshold to the intended and predicted speed. The mean accuracy of the speed detection was 67.89 ± 8.96 % (Matthews correlation coefficient (MCC): 0.35 ± 0.16). By combining the velocity vector into an angle θ of the movement direction, we obtained an average angular error of 88.26 ± 6.79^◦^ as a deviation from the executed movement direction. We also evaluated whether the obtained accuracy in movement detection and direction estimation were significantly better than random performance for each participant by estimating the distribution of a random classifier through simulation. Twelve out of fifteen participants exceeded the level of significant performance for the movement detection and six for the direction. The average levels of significant performance were 59.03 % and 88.34^◦^. Figure 1 F and G show a selection of cursor movements executed during the online target selection task for different targets.

When grouping participants by prior participation in BCI studies (novice: no prior experience, intermediate: 1 − 5 prior participations, experienced: more than 5 prior participations), we observed performances (number of selected targets) of 12.8 ± 4.8 (experienced, 6 participants, p*_s_* = 5.2), 2.0 ± 2.7 (intermediate, 3 participants, p*_s_* = 3.7) and 3.8 ± 3.3 (novice, 6 participants, p*_s_* = 4.3) selected targets. These results, together with the according level of significant performance, are displayed in Figure 1 C. This observed discrepancy in target selection performance between experienced BCI users and novice/intermediate users can similarly be observed in the decoded movements and other measures. The extended groupwise-average results for the performance metrics and the statistics (two-sample t-test) between groups can be found in the supplementary material (Table S1-S5).

### 2.2 Offline Decoding of Continuous Hand Movement

Training and validation data for model calibration was recorded while participants executed 120 continuous hand movements each (Figure 2 A). The executed trajectories of the participants were close to the optimal, intended motions as cued by the calibration paradigm, achieving Pearson correlation coefficients between the executed motion and the intended motion of 0.92 ± 0.02, 0.89 ± 0.03 and 0.87 ± 0.03 for velocities in x- and y-direction and speed, respectively. Figure 2 B shows intended (black) and executed (green) motions of the participants. Since the visual paradigm was the same for all participants, the intended movements were also identical for corresponding trials.

**Figure 2.**
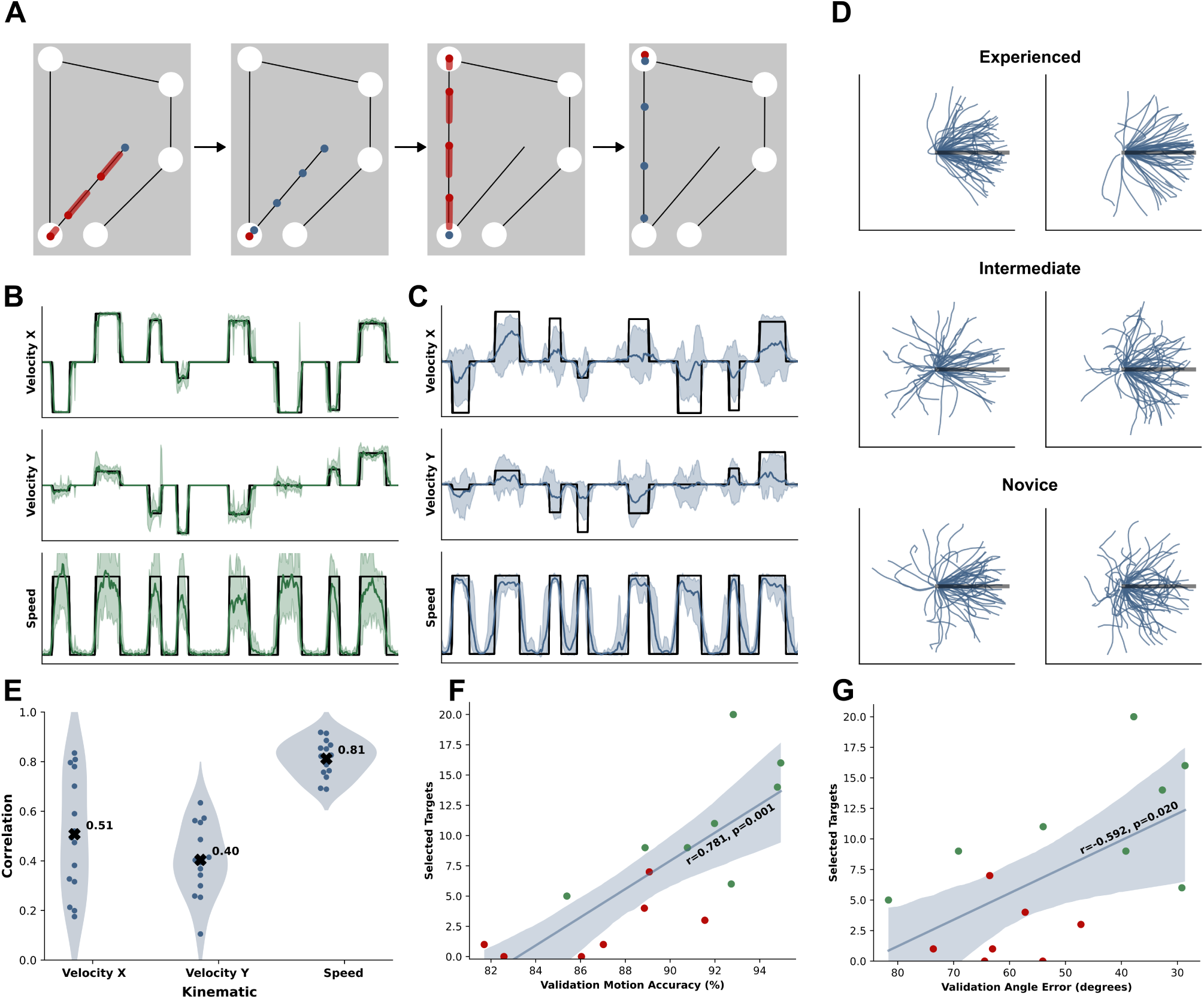
Overview of calibration paradigm and validation results. (A) Calibration paradigm for collecting training data trials. (B) Intended (black) and executed (green) movement showing the velocity in x- and y-direction and the speed. (C) Intended (black) and decoded (blue) movement showing the velocity in x-and y-direction and the speed for the validation set. (D) Cursor movements of three different participants after applying the decoded direction and speed to the recorded neural data (validation set). The motions are normalized an aligned. (E) Participant-wise correlation between the intended and the decoded velocities in x- and y-direction and the speed for the validation set. (F) Correlation between the accuracy of the movement vs. rest classification on the validation set and the online target selection performance. The light-blue area shows the 95 % confidence interval. Datapoints in green achieved a significantly better than chance performance during the online task. (G) Similar to (F), but showing the correlation between the angle error on the validation set and the online target selection performance.

The CNN-LSTM models (individually trained per participant) were trained to predict the speed s (binary, movement vs. rest) and movement direction (velocity vector v⃗ = (v*_x_*, v*_y_*)*^T^*) of the hand. We show these decoded hand kinematics in Figure 2 C in blue together with the intended movement (ground truth). For the validation data, we obtained an average Pearson correlation coefficient between decoded and intended motion of 0.81 ± 0.07 for speed and of 0.51 ± 0.24 and 0.40 ± 0.14 for velocity in x- and y-direction, respectively (Figure 2 E). The Pearson correlation coefficient between decoded and actual movement was slightly reduced at 0.80 ± 0.07 for speed and 0.49 ± 0.24 and 0.37 ± 0.13 for velocity in x- and y-direction, respectively. The average accuracy of the movement vs. rest classification was 89.27 ± 4.10 % (MCC: 0.78 ± 0.08) with an average angular error of 52.51 ± 17.02^◦^. For the predicted versus actual motion, the average accuracy of the movement classification was 88.88 ± 3.95 % (MCC: 0.77 ± 0.08) and the average angular error was 54.67 ± 16.80^◦^. Similar to the online performance, we evaluated whether the obtained accuracy in movement detection and angle error was significantly better than random performance for each participant. On average, the level of significant performance was 54.16 % for the movement detection and 87.53^◦^ for the direction. All participants achieved a performance better than this level, indicating that movement and direction could be identified with a performance that was significantly better than chance.

To improve decoding accuracy, we introduced a decoding delay d during model training by shifting the kinematic data in relation to the neural data. The average decoding delay, optimized per participant, was 0.73 ± 0.17 s, i.e., an executed movement at t = 0.0 s was on average delayed by 0.73 s relative to the neural data.

Figure 2 D shows the normalized (in length and direction) executed and decoded trajectories from the validation set for experienced, intermediate and novice participants. These derived movements correspond directly to the obtained accuracy of the speed detection and direction decoding.

### 2.3 Correlation of Movement and Selection Performance

The correlation between the accuracy of the movement classification during the online task and the selected targets was 0.628 (p = 0.012). The correlation between the angle error of the movement direction (i.e., the error between the angle of the executed motion and the cursor motion) during the online task and the selected targets was −0.480 (p = 0.070). Thus, the accuracy of the movement detection was significantly correlated with the target selection performance, while the directional mismatch between the executed hand movement and the cursor motion, although showing a trend of influencing the number of selected targets, was not significantly correlated with the target selection performance.

We further investigated the correlation of the movement detection accuracy and the angle error of the validation set (calibration paradigm) with the online target performance measures to estimate whether the validation measures were indicative of the online performance. Correlations for motion detection accuracies and angle error (validation vs. online) were 0.658 (p = .008) and 0.556 (p = 0.032), respectively. The correlation between the validation motion detection accuracies and validation angle error with the online target selection performance were 0.781 (p = 0.001) and −0.592 (p = 0.020), respectively. Validation set performances were thus indicative of the performance during the online task. The correlation between the validation motion detection / angle error and the online target selection performance is shown in Figure 2 F and G.

### 2.4 User Perception of Online Cursor Control

Participants were asked to rate their subjective experience regarding the general control, the movement detection performance and the directional accuracy on a visual analog scale. The ratings (Figure 3 A) showed a significantly (paired t-test) higher feeling of control over the speed (rating: 6.0 ± 2.4) than for the direction (rating: 2.2 ± 1.2). The feeling of general control was similarly rated low (rating: 2.5 ± 2.0). There was no significant difference between the directional control and the general control ratings, as assessed by a paired t-test.

**Figure 3.**
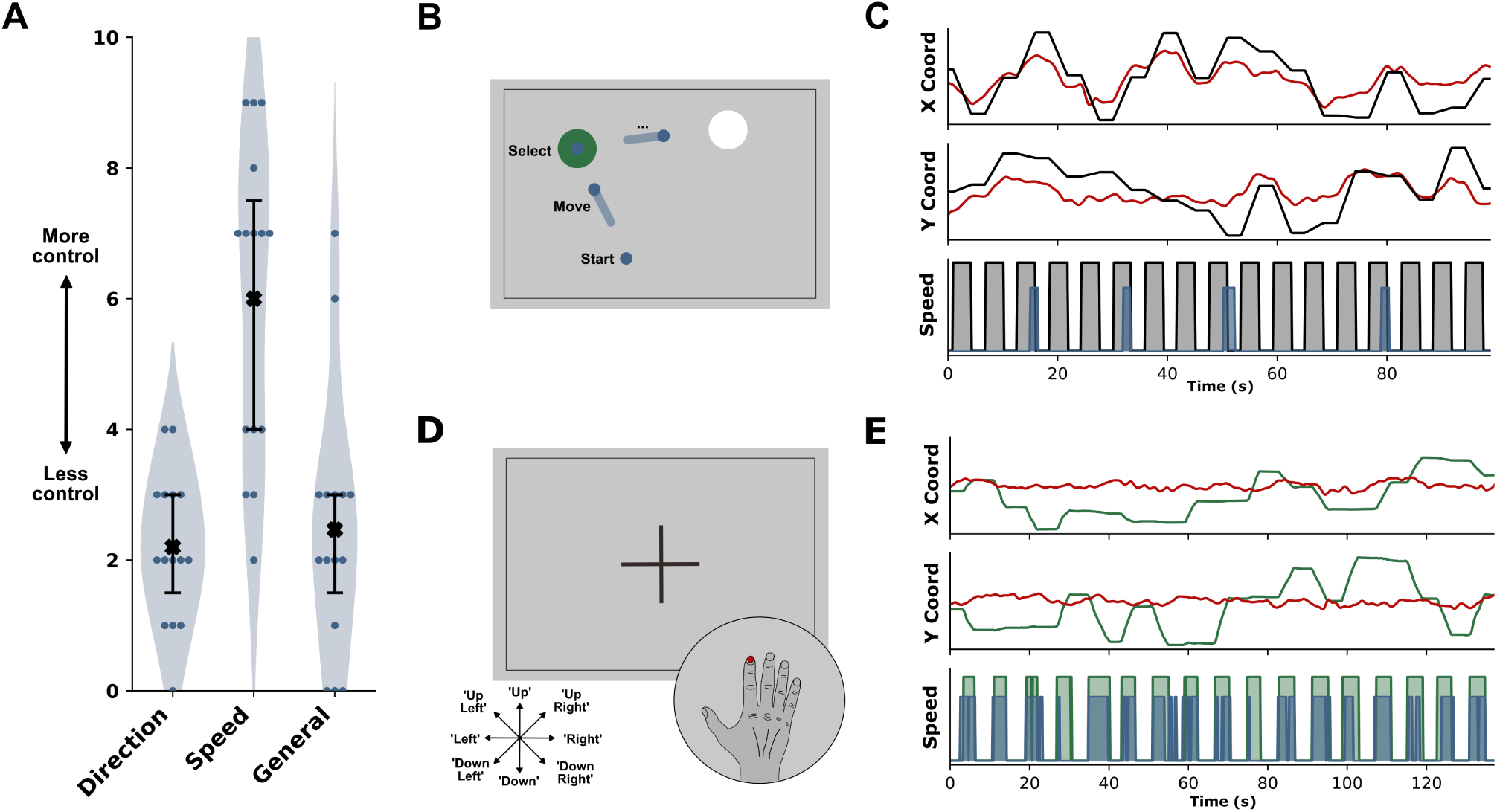
(A) Subjective perception of the participants regarding their general control over the cursor, the speed (movement vs. rest) of the cursor and the direction of the cursor. (B) Observation task intended to evaluate the influence of eye artifacts on the decoded cursor movement. Participants kept their hand fixed while observing a cursor perform an optimal trajectory in the target selection task on the screen. (C) Observed cursor movement speed and motion in x- and y-direction (black), decoded speed (blue) and derived motion in x- and y-direction from the EOG data (red) for an experienced BCI user during the observation task. (D) Auditory movement task intended to evaluate the performance of the decoded cursor movement without the influence of eye artifacts. Participants fixated their gaze on a fixation cross and moved their hand according to auditory commands, while the view of their hand was obstructed. (E) Executed hand movement speed and motion in x- and y-direction (green), decoded speed (blue) and derived motion in x- and y-direction from the EOG data (red) for an experienced BCI user during the auditory movement task.

Participant ratings of control were significantly correlated with the achieved performance in the online target selection task, demonstrating that participants reporting higher levels of control also achieved higher numbers of selected targets during the task. Specifically, the correlation between perceived control over direction, speed and general control over the cursor with the target selection performance were 0.79, 0.87 and 0.84, respectively. The correlation between the perceived control over the speed of the cursor and the accuracy of the movement detection was 0.73 (p = 0.002). The correlation between the perceived control over the direction of the cursor and the angle error was −0.62 (p = 0.013). This indicates that the subjective experience of participants matched the actual performance during the target selection task.

### 2.5 Influence of Eye Artifacts on Decoding Performance

While eye artifacts were removed prior to decoding during the processing pipeline (Kobler et al., 2020a), we additionally conducted two tasks, one auditory movement task and one observation task, aimed at evaluating the influence of eye movements on the decoded trajectory (Figure 3 B and D).

For the auditory movement task in which participants moved their hand according to auditory commands, the Pearson correlation coefficient between the executed motion and the decoded movement showed an average value of 0.60 ± 0.19 for the speed and −0.01 ± 0.09 and 0.00 ± 0.11 for the velocities in x-and y-direction, respectively. Movement detection accuracy and angle error were 81.77 ± 8.14 % (MCC: 0.60 ± 0.19) and 90.07 ± 12.26^◦^ with average levels of significant performances of 56.67 % and 84.05^◦^. All participants obtained detection accuracies better than their individual levels of significant performances for the movement and five participants achieved angle errors below their individual level of significant performance for the angle error. The correlation of the eye movements with the executed hand movements (x- and y-coordinates) was on average −0.04 ± 0.25 and −0.07 ± 0.15.

In the observation task, the decoder output was assessed based on eye movements while the hand remained in a rest position. The correlation between the eye movement and the displayed and observed cursor motion was on average 0.77 ± 0.07 (x) and 0.46 ± 0.14 (y). The average false positive rate (FPR), including periods of cursor rest, was 22.34 ± 26.88 %. In an average of 43.67 % of the observed cursor motions, a decoder movement of more than 0.2 s was detected. For five out of fifteen participants, a movement was detected in less than 10 % during the observed motions. The Pearson correlation coefficient of the decoded direction with the displayed cursor direction was −0.03 ± 0.15 and 0.01 ± 0.10 for velocities in x- and y-direction and 0.10 ± 0.10 for the movement speed.

Selected hand movements, eye and decoder trajectories for both tasks are shown in Figure 3 C and E.

### 2.6 Neural Correlates of Cursor Control

We investigated neural correlates of speed and direction during continuous hand movement by obtaining the feature importance of the model for frequency-specific channels and brain areas and mapped the feature importance of the frequency-specific channels onto the cortex of the brain using source localization. The grand average of the individual participants’ feature importance maps for the frequency ranges 0.3 − 2.0 Hz, 8 − 14 Hz, 25 − 30 Hz are shown in Figure 4 A. The feature importance for the overall frequency range is shown in Figure 4 B. A comprehensive analysis of all frequency ranges in narrow bands is given in the supplementary material (Figure S1). Positive values (in arbitrary units, a.u.) indicate a higher importance of the brain area and frequency band to the decoder. For reference, Figure 4 C comprises an anatomical overview of the brain areas (frontal, parietal, occipital and temporal lobe). High feature importance for the speed was localized mainly around contralateral areas of the sensorimotor cortex (M1 and S1) in the 25 − 30 Hz frequency range, which corresponds to high-beta frequencies. Specifically, the hand area of the sensorimotor cortex corresponded well with the important brain areas. For the directional feature importance, parieto-occipital areas in the 8 − 14 Hz range (alpha frequency range) and contralateral central motor areas in the 25 − 30 Hz range showed the highest importance. For the directional decoding, brain areas in parieto-occipital and contralateral motor areas in the delta frequency range (0.3 − 2 Hz) showed increased feature importance, although the relevance was reduced compared to higher frequency ranges.

**Figure 4.**
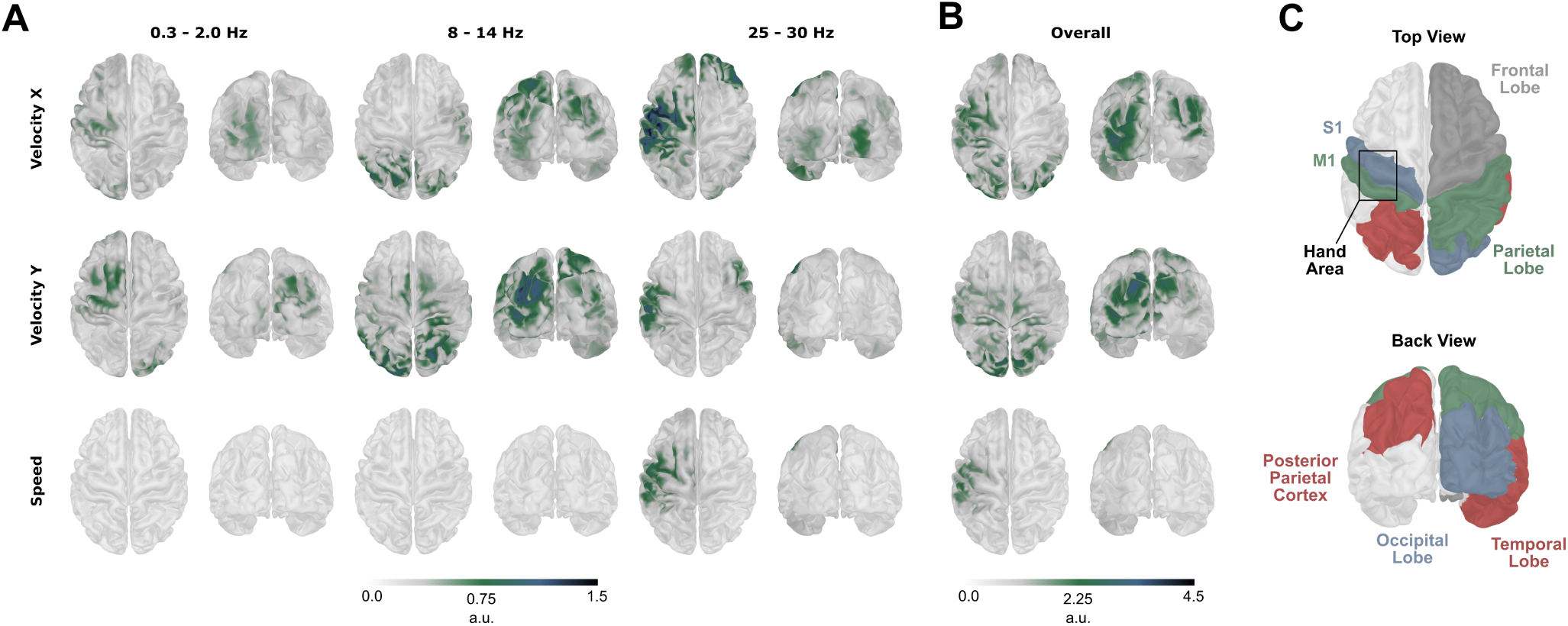
(A) Feature importance of brain areas in specific frequency bands in arbitrary units (a.u.) based on the difference compared to original performance. Higher values indicate a higher importance of the brain region to the model. (B) Feature importance of brain areas over all frequencies in arbitrary units (a.u.) based on the difference compared to original performance. (C) Anatomical parcellation of the brain according to (Van Essen, 2005; Desikan et al., 2006) with areas related to hand movement and coordination. Top view and back view correspond to the views shown in (A) and (B).

## 3 DISCUSSION

The presented continuous cursor control framework demonstrates sufficient precision to enable target selection in a substantial number of participants. Compared to prior EEG-based BCIs for cursor control (Mondini et al., 2020; Lu et al., 2026; Forenzo et al., 2024), our study demonstrates continuous control based on free hand movements not aligned to fixed motions (e.g., pursuit tracking tasks) and allows for no-movement states between continuous motions. Overall, the study shows the feasibility of the proposed data collection and decoder training procedure for application in an online environment and indicates the practicality of the approach in real-world scenarios.

While invasive BCIs routinely enable high target selection accuracies of over 90 % (Patrick-Krueger et al., 2024), the achieved target selection rate of 17.7 % overall and 32.1 % for experienced users in this study is substantial for a non-invasive, EEG-based BCI. Comparisons between previous BCIs for cursor-controlled target selection are difficult due to differences in the task implementation such as time-outs, dwell-period or boundary behavior. Further and to our knowledge, no EEG-based (non-invasive) BCIs exist that enable the achieved form of free, continuous motion from hand movement and real-world applicability as demonstrated with the target selection task. In our study, target selection was enabled by the introduction of a no-movement state, which is essential for natural control and real-world applications. The introduction of the start-and-stop capabilities in the current framework is therefore an important contribution to the state of the research in this field.

Specifically, while existing studies controlling a cursor based on modulation of neural signals demonstrated sufficient control of the cursor movement (Wolpaw and McFarland, 2004), they lack the intuitive control achieved from direct utilization of hand movements and require an extensive learning phase. Additionally, such control methods do not enable idle no-movement periods or dwell-time selection.

The substantial number of selected targets during the online target selection task, especially for the group of experienced users, shows that participants could effectively utilize the framework to execute planned cursor movements. The variation in the number of selected targets between groups of experienced and inexperienced users indicates the importance of familiarity of users with BCI-controlled interfaces. We consequently found an improvement of the ratio of successful target selections with time during the online task, further indicating a dependence of the performance on the familiarity with the task and interface. However, the increased correlation of the decoded hand kinematics found in the validation data also points towards differences in the collected training data between experienced users and novices which cannot be explained by familiarity with BCI-controlled interfaces. In line with this observation, previous studies have found a positive influence of user training on BCI performance (Abu-Rmileh et al., 2019) and cortical activations related to movement (Kaiser et al., 2014). Overall, this indicates that the achievable performance of the proposed framework is close to the performance of experienced users in this study.

Our framework achieved overall higher performance than previous studies for the decoding of kinematics from continuous hand movements. The correlation of the decoded velocities in x- and y-direction on the validation data from the calibration task exceeded previously obtained values of around ∼ 0.45 (v*_x_*) and ∼ 0.35 (v*_y_*) (population averages) (Borra et al., 2023) or lower (Lv et al., 2010; Bradberry et al., 2010). In accordance with these results, we similarly found reduced correlation for velocities in y-direction compared to those in x-direction. Similar reduced correlation in y-direction was also observed for the position instead of the velocities in related work (Mondini et al., 2020). Previous studies on the detection of continuous movement vs. rest activity reported lower accuracies than our achieved results with recent studies demonstrating an MCC of ∼ 0.6 (Crell et al., 2026). The high correlation of the speed and accuracy of the movement vs. rest detection is an important finding of our study. Nearly all participants obtained significantly better than random accuracies for the movement detection and, overall, ratings of control over the movement speed were high. Participants also verbally reported a feeling of confidence that ”the cursor moved when they wanted it to move” during the online task.

The reduced correlation between the executed and decoded speed in the online task can mostly be attributed to the reduction in uniform motions. During the guided motions in the calibration task, visual feedback was directly given to the participants and their movement speed was kept at a reduced, nearly constant rate. Since participants could move freely during the online task, motions tended to be more diverse in the executed speed and duration and thus lead to reduced correlation with the decoder output. During the auditory movement task, where participants were clearly instructed to move for a certain duration at a constant pace, the obtained correlation of the speed was close to the performance on the validation set.

While the movement detection successfully allowed start-and-stop movements with variable durations during the online task, the correlation of the decoded movement direction with the executed movement dropped below the level of significant performance in most participants. This behavior did not improve during more controlled motions in the auditory movement task. While directional correlation for experienced BCI users remained slightly higher than those of less experienced participants, the obtained angular error indicates near-random performance for the movement direction. This is also indicated by low subjective ratings of control over the cursor movement direction. In previous studies of online cursor control based on pursuit tracking tasks with similar calibration and online tasks, no such performance drops were reported (Mondini et al., 2020). We therefore assume that the change in directional decoding capabilities stems from differences in the neural patterns between the guided (calibration) and free (target selection) online task. The selection of targets despite the missing directional control may be attributed to start-and-stop motions with differing duration: if participants started a movement directed at the target, they could extend the motion, while they would immediately stop the movement if the direction was wrong. This is also emphasized by the significant correlation of speed and target selection performance compared to the non-significant correlation of the number of selected targets with the angle error. More broadly, this observation has important implications for the BCI field, as many studies evaluate movement decoding exclusively in offline settings using visually guided paradigms. In such cases, offline performance may overestimate the degree to which the decoder captures neural correlates of movement itself and may not accurately predict performance during self-paced online control. The marked discrepancy between offline and online directional decoding observed in the present study highlights the importance of validating continuous-control BCIs in realistic online environments.

Results from the feature importance analysis indicate the utilization of different neural correlates for velocities and speed by the model. For the decoding of speed, mainly information from contralateral central channels around C3 in the high-beta band was utilized. The focus on contralateral motor areas is in accordance with literature (Wang et al., 2022; Yuan et al., 2010; Crell et al., 2026; Kobler et al., 2020b) and indicates an emphasis on event-related (de-)synchronization patterns occurring during continuous movement (Pfurtscheller and Silva, 1999). Velocities in both x- and y-direction also showed dependence on high-beta frequencies in central, contralateral motor areas. Further relevant information was located in alpha frequencies at parieto-occipital areas in the posterior-parietal cortex. Parieto-occipital and motor areas have previously been identified as key structures during visually guided online control of hand motions (Ferraina et al., 2009), with specifically areas at the parieto-occipital junction being tuned to the combination of visual target location with hand position and the direction of the motion (Battaglia-Mayer, 2019; Battaglia-Mayer and Caminiti, 2019). Similar activation patterns for directional activity have been identified in previous EEG studies (Mondini et al., 2020; Kobler et al., 2020b) with one particular study identifying similar high-beta frequency components relevant to velocity decoding (Lv et al., 2010). However, these studies also found parieto-occipital areas being involved during observation of cursor movements. Main differences between execution and observation were found in the involvement of the motor areas (Kobler et al., 2020b). This finding could indicate observation-specific neural patterns as the reason for the observed performance drop between visually guided motions in the observation task and free or speech-guided motions in the online and auditory movement task. We assume that the calibration task could have induced neural correlates related more to the visual distance of the cursor from the displayed guidance line than to the target.

While visual input may contribute to performance differences, our preprocessing and model-training pipeline was specifically designed to minimize the influence of eye movement–related activity from the EEG signals. Feature importance analysis also indicates little dependence on frontal channels, suggesting that eye movement–related activity did not substantially contribute to model performance. Further, we introduced the auditory movement and observation tasks to provide a quantization of the (in-)dependence of the decoded movements on eye artifacts and eye motions. Results from these tasks showed clear independence of the decoded speed from eye movements as well as a lack of correlation between the decoded motion and the movement of the eyes.

Our proposed cursor control framework and online application demonstrates the importance of no-movement states for a successful target selection. Future work should focus on the transfer of directional decoding performance results achieved during visual, guided motions to free, unguided cursor movements. Additionally, while the current study was conducted with able-bodied participants and executed hand movements, the system is directly applicable to motor-impaired individuals. Further investigations should be conducted with the direct involvement of the target group to identify whether similar performances can be achieved and finally apply the framework in a real-world setting.

## 4 METHODS

### 4.1 Participants and Setup

Fifteen participants (10 females and 5 males with an average (± standard deviation) age of 26.7 ± 3.5 years) took part in the study. EEG data of each participant was obtained from 60 channels. The EEG setup consisted of a standard 10-10 setup with additional electrodes at PPO1h and PPO2h. Channels at Fp1, Fpz, Fp2, F9, F10, FT9, FT10, T9, T10, TP9, TP10, P9 and P10 were excluded. Four electrooculography (EOG) electrodes were positioned on the outer canthi of the eyes and above and below the left eye. EEG and EOG data was recorded with biosignal amplifiers (BrainAmp, Brain Products GmbH, Gilching, Germany) at a rate of 500 Hz.

Motion capture data (coordinates along x- and y-axes along the table orientation) of the right index finger was obtained with a custom marker-based tracking system mounted above the hand of the participant. The tracked area (34 × 63 cm) was covered in a friction-reducing surface. Participants were positioned 100 cm in front of a screen with the chair height adjusted to fit the height of the eyes with the center of the screen.

The study was conducted in accordance with the Declaration of Helsinki and was approved by the Ethics Committee at Graz University of Technology (EK-49/2024). Participants received monetary compensation for their time and provided written informed consent before study participation.

### 4.2 Calibration Paradigm

During the calibration paradigm (Figure 2 A), participants controlled a blue cursor (participant cursor) on the screen with their right hand. Movements of the hand were tracked with a motion capture system and mapped to the cursor while applying a constant speed. Specifically, the participant cursor was controlled using a fixed speed, regardless of the actual speed of the hand movement, and the direction of the movement in the form of the velocity in x- and y-direction. Participants were therefore only able to control the direction and the timing of the cursor movement, but not the speed at which the cursor moved. This behavior was chosen since it ensured a high accuracy of the executed movement with the movement intended from the visual paradigm. Intended movements of the participant cursor were cued by a displayed trajectory and a red cursor (paradigm cursor). This cursor had an additional tail comprising the last 0.8 s of the movement. Trajectories comprised 3 to 7 targets (four trials per number of targets) indicated by white circles and black lines between the targets. The start position of each trajectory was in the middle of the screen.

At the start of each trial, participants positioned their right index finger at the center of the tracked area. Paradigm and participant cursors were reset to the center of the screen. After 2.0 s, the paradigm cursor moved from the starting point to the first target. As soon as the cursor reached the target and the tail of the cursor disappeared, participants moved their cursor along the black line to the target. After a variable waiting period, the paradigm cursor would then move towards the next target and cue the next participant movement. After the last target, the next trial would be displayed and the cursors were reset to the starting position.

The complete task comprised twelve runs of ten trials each. Trajectories had an average duration of 47.6 s. Participants could take breaks after each run, but had a mandatory break after the sixth run to reduce (unnoticed) fatigue.

Prior to the calibration task, participants executed an eye-movement paradigm comprising specified eye movements. The eye movements consisted of rest, horizontal and vertical eye movements and blinks in accordance with stimuli on the screen (Kobler et al., 2020a). The complete paradigm had an approximate duration of 6.3 min.

### 4.3 Online Task

The online target selection task comprised eight runs of five targets each and had an average duration of 21.6 min. Participants controlled the speed and direction of a blue cursor through decoded EEG and had to select targets on the screen using this cursor. A target could be selected by positioning the cursor within the target for 3.0 s. The required duration of 3 s was indicated by a green progress circle within the target (Figure 1 B). While the cursor was positioned within a target, the target gradually filled with the progress circle. The circle expanded from the center of the target to visually indicate the progress of the dwell-time selection to the participants. If the cursor left the target, the progress circle would recede again at the same rate as when growing. The shrinking of the progress circle would start only 0.2 s after the cursor left the target. If the cursor re-entered the target, the progress circle would start to grow from its current expansion in case it was not completely shrunk. The target was selected when the progress circle completely filled the target. When a target was selected, the next target was immediately displayed. The next target was also displayed when a time-out of 30 s was exceeded. Targets were randomly generated at a fixed distance from the cursor position at the time of the target generation.

At each new run, participants started at the center of the screen. However, the cursor position was not reset after a target selection or time-out. If participants reached the outer bounds of the screen, the cursor was mapped to the corresponding opposite side of the screen.

### 4.4 Auditory Movement and Observation Tasks

Eye movements are a major source of artifacts in EEG measurements. Since not only eye blinks but also rotations of the eye when following a visual object generate electrical artifacts with a high amplitude that can be measured directly, especially in frontal channels, the issue of decoding movement instead of gaze needs to be carefully addressed. To this extent, two specific tasks were introduced, the auditory movement and the observation task. During both tasks, participants did not control a cursor.

In the auditory movement task (Figure 3 B), only a fixation cross was shown. Participants were instructed to focus the fixation cross with their gaze and move their hand according to auditory commands given from a speaker. These commands comprised *up*, *down*, *left*, *right* and the diagonal movements *up-left*, *up-right*, *down-left* and *down-right*. On hearing a command, participants moved their hand in the corresponding direction for approximately 2.0 s and then waited for the next command. Commands were issued every 6.0 s with 20 commands in total.

The observation task (Figure 3 C) was visually similar to the online task. In this task, the cursor was not controlled by the participant but instead moved automatically along the optimal trajectory to select the targets. In total, 20 targets were displayed and selected. Participants were instructed to keep their hand at the resting position and follow the moving cursor with their eyes.

### 4.5 Data Processing

EEG was first bandpass-filtered between 0.3 and 70 Hz (Butterworth filter, 4*^th^* order) and powerline noise was removed using a 50 Hz notch filter. Eye artifacts were eliminated with a correction matrix derived from the eye-movement data collected prior to the calibration paradigm (Kobler et al., 2020a). EEG was then lowpass-filtered at a cutoff frequency of 40 Hz and re-referenced to the common average of all EEG channels. The data was then subsampled to a sampling rate of 125 Hz.

Online and offline data was processed using BOLT (Brain OutLeT), a python-based framework for high-performance, real-time deployment of neural decoding models and a proprietary extension of ezmsg (https://github.com/ezmsg-org/ezmsg) developed by ABILITY Neurotech, Geneva, Switzerland.

### 4.6 Decoder and Calibration

To obtain values of the movement vector, we trained a model to predict the amplitude (speed) and the direction (velocity in x- and y-direction) from windows of the neural data. We utilized a CNN-LSTM model based on the EEGNet (Lawhern et al., 2018) architecture. The detailed model architecture can be found in the supplementary material (Table S6). Previous research suggested that the combination of CNN and LSTM enables higher performances than single CNNs for movement tasks (Chen et al., 2022; Lu et al., 2026). Conceptually, the EEGNet architecture extracts relevant spatial and spectral features while the LSTM layer introduces temporal dependency on the previous state of the model output. This is especially relevant in the case of continuous movements, since the direction of each step during a continuous motion is, to some extent, dependent on the direction of the previous step. To further ensure smooth subsequent predictions of the model, we applied a custom loss function (sMSE, *smooth mean-squared error*) during training, comprising the mean squared error (MSE) and an additional *smoothness* term penalizing large differences between consecutive model predictions (Equation 1).

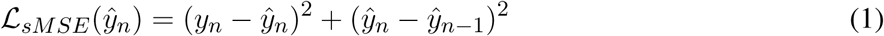

We trained the model on ten out of twelve runs of the calibration paradigm and utilized the remaining two runs for validation and hyperparameter selection. We extracted training and validation data as windows of the pre-processed EEG. Windows were obtained with a variable window length *wl* ∈ [2; 3]s as a hyperparameter. For each window, we extracted the label from the intended or executed movement. Specifically, we extracted movement speed and velocities in x- and y-direction by calculating the derivative of the x- and y-coordinates. The speed (i.e., the magnitude of the velocity vector 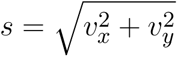 at each time step was normalized to a length of one at all points where the magnitude exceeded a fixed threshold. At all other time points, the speed and velocities were set to zero. Velocities were similarly normalized in accordance with the speed, so that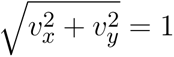 . Data windows were labeled with the binary movement speed and the normalized velocities.

We further employed the label delay d as an additional hyperparameter. Based on previous research showing that the optimal classification performance can be achieved by delaying the classification relative to the movement onset and incorporate further EEG samples (Ofner et al., 2019; Crell et al., 2025, 2026), we optimized the delay in the range d ∈ [0.25, 0.5, 0.75, 1.0]s. This delay constitutes a decoding delay between the label and the decoded motion.

For each participant and each window length wl and delay d combination, an individual model was trained. In order to enable usage of the framework also for motor-impaired people, for which only attempted movement can be utilized, we inferred the motion of the participants from the visual calibration paradigm. Specifically, the visual cues of the calibration paradigm, in the form of the movements of the paradigm cursor, were given such that the executed motions of the hand could be inferred directly and accurately from the paradigm. These motions inferred from the paradigm, called *intended motions*, were then utilized for training of the model, rather than the executed motions. We employed early stopping based on validation loss with a patience of ten epochs. The correlation of speed and velocities on the validation set was then calculated by obtaining the consecutive model output of the ordered windows in each trial of the validation set. We calculated the trialwise Pearson correlation coefficient for each kinematic between the model output and the intended movement. Hyperparameters wl and d were selected based on the highest average correlation of the three decoded kinematics.

To obtain binary movement vs. rest classification, we further applied an optimal threshold τ to the speed. The threshold was selected by optimizing the accuracy of the movement vs. rest classification on the validation set data of the best performing model.

### 4.7 Offline Evaluation

We calculated the trialwise Pearson correlation coefficient between the decoded and the intended speed and velocity in x- and y-direction on the validation set. We additionally calculated the trialwise Pearson correlation coefficient between the executed and the decoded motions. We further obtained the accuracy and the Matthew’s Correlation Coefficient (MCC) for the detection of movement vs. rest after thresholding the decoded speed. The MCC was chosen since it is robust to class imbalance between the rest and movement class (Chicco and Jurman, 2020). The directional accuracy was assessed by obtaining the direction from the decoded velocity in x- and y-direction and calculating the angular error between the decoded and the intended/actual angle of the movement vector. The angular error (in degree) was wrapped at ±180^◦^ to account for circular properties of the angle.

We further obtained subject-wise levels of significant performance for the accuracy of the movement detection and the angle error. This was achieved by estimating the distribution of a random classifier through simulation (n = 10000) and utilizing the 95*^th^* percentile of this distribution as a level of significant performance (Müller-Putz et al., 2008).

### 4.8 Online Target Selection Task Evaluation

The Pearson correlation coefficients, MCC, movement accuracy and angular error for the decoded movement during the online target selection task were calculated similar to the metrics during the calibration paradigm. All metrics were calculated between the decoded kinematics and the kinematics derived from the executed movement. Trialwise correlation was calculated between the appearance of a target and the disappearance of this target in the online target selection task.

Targets were deemed correctly selected if the complete target circle was filled out by the progress circle. While a time-out of 30 s was applied to the visualization of the circles, targets would also be deemed correctly selected if they were selected within 0.5 s after the timeout.

The statistical evaluation of the number of selected targets was based on a conditional randomization test of the executed cursor movements of each participant (Crell and Müller-Putz, 2026). This method was applied to evaluate whether the observed number of selected targets was significantly (α = 0.05) better than random performance. For this, we generated random targets that were not overlapping with the actual target for each trial and evaluated the number of random targets that were selected based on the simulated dwell-time. We then obtained the null distribution of random performance by repeating this procedure for 1000 times. We obtained the 1 − α percentile of the null distribution above which performance was considered significantly better than chance for each individual participant.

We identified trends over runs for the target selection performance by using a population-averaged generalized estimating equations (GEE) logistic model (Liang and Zeger, 1986). GEE was selected since it can analyze correlated binary outcomes from repeated measurements within subjects. The target selection performance was modeled as a function of the run number, with clustering by subject to account for repeated measures. Results are reported as odds ratios, which were obtained by exponentiating the model coefficients. We also examined changes in time-to-target selection for successful trials. Times were log-transformed to reduce skewness. A linear mixed-effects model (Lindstrom and Bates, 1988) was fitted with run number as a fixed effect and subject as a random intercept.

Cross-task correlations between validation set performances and target selection performance were assessed by calculating the Pearson correlation coefficient. All statistical tests were performed under a significance level of α = 0.05

### 4.9 Behavioral Analysis

Participants were asked to evaluate their perceived control over the cursor after completing the online target selection task. The questionnaire comprised three visual analog scales on which participants rated their control over the cursor speed (movement vs. rest), the movement direction and the general feeling of control over the cursor. Each scale ranged from zero (no control) to ten (complete control). We tested for statistically significant differences between the perceived control of the speed and the direction with a paired t-test. Ratings were further evaluated regarding their correlation with the target selection performance based on the Pearson correlation coefficient. We also evaluated the correlation between the perceived control of the speed and the direction with the accuracy of the movement detection and the angle error, respectively. All statistical tests were performed under a significance level of α = 0.05.

### 4.10 Auditory Movement and Observation Task Evaluation

Horizontal (x) and vertical (y) eye movements were extracted during both tasks from the EOG data by lowpass-filtering the data above 4 Hz (Butterworth, 4*^th^* order) and removing linear trends to eliminate low-frequency shifts. We then obtained the equivalent of horizontal coordinates by subtracting the EOG channel next to the right canthus of the eye from the EOG channel next to the left canthus. Similarly, we obtained the equivalent of vertical coordinates by subtracting the EOG channel below the eye from the EOG channel above the eye.

We measured the Pearson correlation coefficient between these eye coordinates and either the executed hand movement or the observed cursor movement. Apart from the correlation, we also evaluated similar measures as for the online and calibration paradigms.

### 4.11 Analysis of Neural Correlates

Neural correlates of movement and direction decoding were analyzed by investigating the frequency-specific channel-wise feature importance. The method to obtain the feature importance has previously been successfully applied to the investigation of feature importance for the detection of hand movements (Crell et al., 2026; Kostoglou et al., 2016). Here, we further extended this investigation to relevant frequency bands.

For the channel-wise feature importance, we utilized the validation set from the calibration paradigm and eliminated specific frequency content from individual channels by applying a band-stop filter (Butterworth, 4*^th^* order). We applied band-stop filters with a width of 2 Hz and in steps of 2 Hz between 0.0 and 20 Hz and in steps of 5 Hz between 20 and 40 Hz. The feature importance was assessed individually per participant. For each channel c and frequency band f, a band-stop filter was applied and the performance was evaluated on the adapted validation data. We measured the change in performance Δp*_c,f_* (correlation for speed and velocity in x- and y-direction) for each channel and frequency band as percentage from the original value. Grand average feature importance was calculated from the average of all participants. The overall channel importance was obtained as the sum of the individual frequency-band specific feature importance values. We investigated the feature importance for multiple narrow frequency bands (see Figure S1 in the supplementary material). Based on results for these narrow frequency bands, frequency ranges with similar patterns were identified and grouped together.

To identify relevant brain areas corresponding to the feature importance of the channels, we mapped the frequency-specific and overall feature importance onto the cortex of the brain using source localization. Source localization was performed using MNE-Python (Gramfort et al., 2013). A cortically constrained distributed source model was constructed on the standard brain template. Source reconstruction was performed using standardized low-resolution electromagnetic tomography (sLORETA), a distributed inverse solution method that yields smooth, depth-weighted current density estimates with reduced localization bias (Pascual-Marqui, 2002).

## DATA AVAILABILITY STATEMENT

Data will be made available upon reasonable request.

## ACKNOWLEDGMENTS

The authors thank all participants for their contribution to this study. This work was funded by the European Union’s HORIZON-EIC-2021-PATHFINDER CHALLENGES program under grant agreement No 101070939 and by the Swiss State Secretariat for Education, Research and Innovation (SERI) under contract number 22.00198. This study was supported by TU Graz Open Access Publishing Fund.

## AUTHOR CONTRIBUTIONS

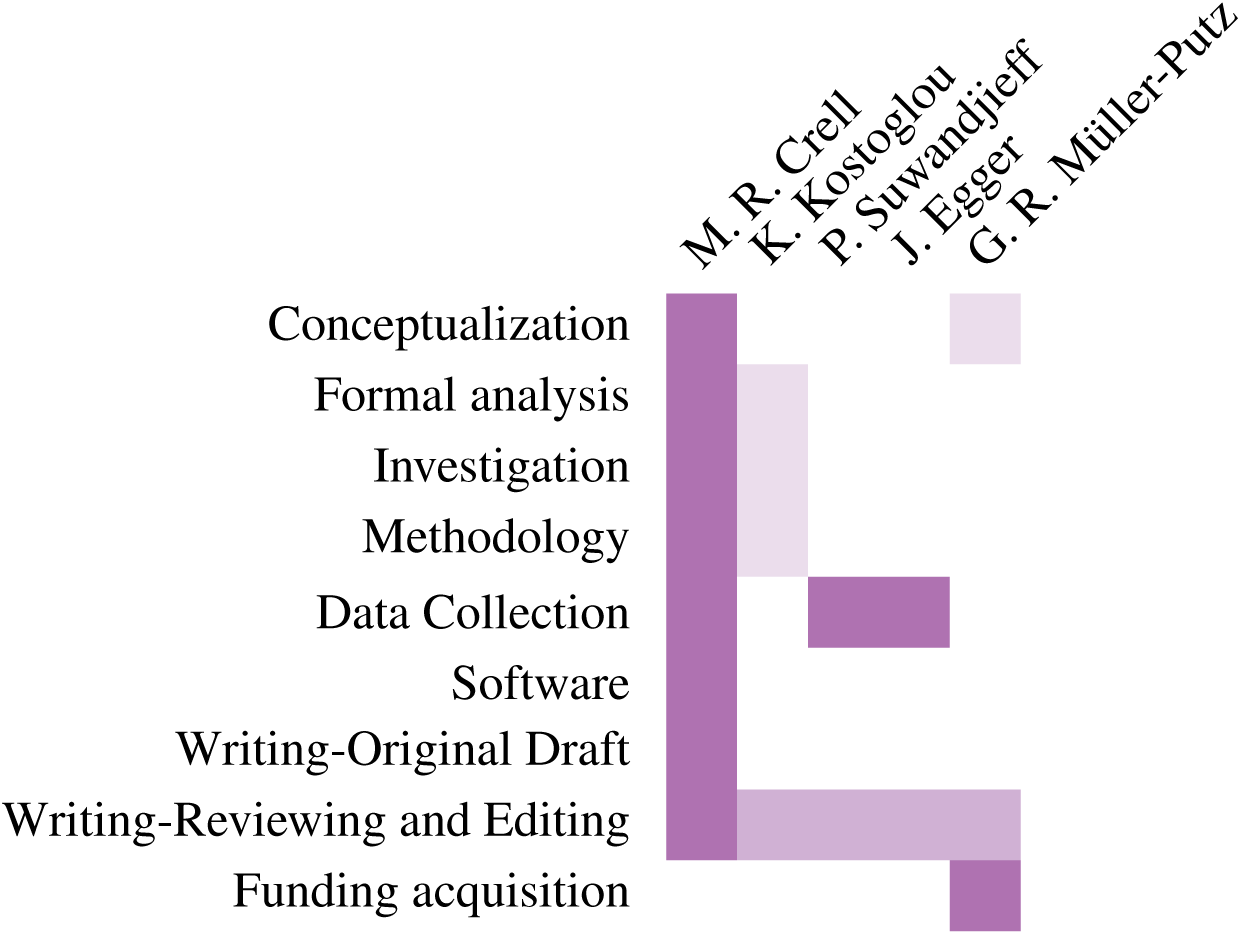

